# Functional interaction of electrical coupling and H-current and its putative impact on inhibitory transmission

**DOI:** 10.1101/2025.02.18.638878

**Authors:** Federico F. Trigo, Pepe Alcamí, Sebastian Curti

## Abstract

The flow of information within neural circuits depends on the communication between neurons, primarily taking place at chemical and electrical synapses. The coexistence of these two modalities of synaptic transmission and their dynamical interaction with voltage-gated membrane conductances enables a rich repertoire of complex functional operations. One such operation, coincidence detection, allows electrically coupled neurons to respond more strongly to simultaneous synaptic inputs than to temporally dispersed ones. Using the mesencephalic trigeminal (MesV) nucleus—a structure composed of large, somatically coupled neurons—as an experimental model, we first demonstrate that electrical coupling strength in the hyperpolarized voltage range is highly time-dependent due to the involvement of the IH current. We then show how this property influences the coincidence detection of hyperpolarizing signals. Specifically, simultaneous hyperpolarizing inputs induce larger membrane potential changes, resulting in stronger IH current activation. This, in turn, shortens the temporal window for coincidence detection. We propose that this phenomenon may be crucial for networks dynamics in circuits of electrically coupled neurons that receive inhibitory synaptic inputs and express the IH current. In particular, molecular layer interneurons (MLIs) of the cerebellar cortex provide an ideal model for studying coincidence detection of inhibitory synaptic inputs, and how this operation is shaped by the voltage-dependent conductances like the IH current, potentially impacting on motor coordination and learning.

## Introduction

Neural circuits are the substrate of information processing by nervous systems, and their output is critical for functions like sensory perception, organization of behavior and cognitive processes. It has long been recognized that the dynamics of neural circuits rely on the interplay between synaptic and intrinsic neuronal properties. Intrinsic properties refer to the characteristics of individual neurons, arising from the repertoire, density and distribution of ion channels in the neuronal membrane, as well as the neuron’s morphological characteristics (Llinas, 1988). Equally essential to circuit function are synaptic properties, which encompass the modality of synaptic transmission (chemical or electrical), the sign of synaptic actions (excitatory or inhibitory), synaptic strength, time course, as well as the network’s connectivity pattern (Getting, 1989). Chemical transmission involves the release of a chemical signal, the neurotransmitter, whose action is typically restricted to the immediate postsynaptic cell in the case of classic neurotransmitters like GABA or glutamate, or to more distant targets in the case of volume transmission. Conversely, electrical synaptic transmission is a communication modality based on the direct flow of electrical currents between neurons through low-resistance pathways consisting of intercellular channel aggregates known as gap junctions.

Extensive experimental evidence indicates that both modalities of synaptic transmission coexist in the vertebrate brain (Martin and Pilar, 1963; Lin and Faber, 1988; Bartos et al., 2002; Galarreta and Hestrin, 2002; Deleuze and Huguenard, 2006; Alcami and Marty, 2013; Alcami et al., 2021). Furthermore, their dynamic interaction with intrinsic neuronal properties underlies essential functional operations that significantly enhance neural networks information processing (Llinas, 1988; Bennett and Zukin, 2004; Connors and Long, 2004; Pereda et al., 2013; Alcamí and Pereda, 2019; Curti et al., 2022). Among these operations, coincidence detection allows neuronal circuits to prioritize synaptic inputs based on their temporal patterns. In fact, while temporally distributed inputs to electrically coupled neurons can be relatively inefficient, synchronous synaptic inputs tend to produce a more reliable activation of postsynaptic neurons (Chillemi et al., 2007; Hjorth et al., 2009; Rabinowitch et al., 2013).

Whereas coincidence detection in circuits of electrically coupled neurons has been extensively characterized both experimentally and theoretically for excitatory inputs, much less attention has been given to coincidence detection in the context of inhibitory synaptic transmission. This phenomenon may be particularly relevant in electrically coupled networks receiving inhibitory synaptic inputs, especially in networks of inhibitory interneurons, which are commonly interconnected through both electrical and chemical inhibitory synapses. Such networks can be found in various brain regions including the neocortex, hippocampus, thalamus, and cerebellar cortex (Mann-Metzer and Yarom, 1999; Bartos et al., 2002; Hestrin and Galarreta, 2005; Deleuze and Huguenard, 2006), where they play a crucial role in modulating principal neuron activity and, consequently, the output of these brain structures.

In this article, we first review the interaction between electrical synapses and intrinsic neuronal properties, highlighting their role in coincidence detection within networks of electrically coupled neurons. Then, based on the novel analysis of data from neurons of the mesencephalic trigeminal (MesV) nucleus, we explore the contribution of active membrane currents to the phenomenon of coincidence detection of hyperpolarizing signals. MesV neurons are electrically coupled, and their electrophysiological properties are well characterized (Khakh and Henderson, 1998; Pedroarena et al., 1999; Wu et al., 2001; Enomoto et al., 2006). In a previous study, we reported increased coupling strength at the peak of hyperpolarizing responses compared to the steady state (Curti et al., 2012). Here, we demonstrate that the hyperpolarization-activated (IH) current confers specific time-dependent properties to electrical coupling strength and coincidence detection. However, since MesV neurons do not receive synaptic hyperpolarizing inputs (Hayar et al., 1997; Verdier et al., 2003, 2004), the physiological significance of hyperpolarizing signals in these cells remains speculative. Nevertheless, we hypothesize that the IH current may endow neural networks with unique computational capabilities by modulating the detection of coincident inhibitory synaptic inputs in circuits of coupled neurons, including interneuron networks. We propose that this effect may be particularly relevant in shaping network dynamics of cerebellar local inhibitory circuits.

### Impact of neuronal intrinsic electrophysiological properties on electrical synaptic transmission

The efficacy of transmission through electrical synapses depends primarily on the gap junction’s resistance, which is determined by the number of intercellular channels, their unitary conductance and open probability (Harris, 2018). The passive electrophysiological properties of the postsynaptic neuron, particularly its input resistance (Rin) and time constant, are also critical factors that influence transmission efficacy (Bennett, 1966). In addition, active membrane conductances can modify coupling strength (Pereda et al., 2013; Curti et al., 2022). For example, the persistent Na⁺ current, a fast inward current that activates at subthreshold membrane potentials, can amplify coupling potentials, thereby enhancing firing probability and promoting synchronous activity among coupled neurons (Mann-Metzer and Yarom, 1999; Curti and Pereda, 2004; Dugué et al., 2009; Curti et al., 2012). On the other hand, K^+^ currents such as I_D_ or I_A_ might also regulate the ability of presynaptic active neurons to recruit their coupled cells (Dapino et al., 2023), and even in combination with the passive membrane properties impart band-pass properties to electrical synaptic transmission instead of the typical low-pass filter characteristics (Hutcheon and Yarom, 2000; Curti et al., 2012; Pereda et al., 2013; Alcamí and Pereda, 2019). Thus, intrinsic electrophysiological properties are capable of fine-tuning neuronal communication through gap junctions in order to match the frequency content of biologically-relevant signals (Curti et al., 2012; Pereda et al., 2013; Davoine et al., 2020).

### Coincidence detection in circuits of electrically coupled neurons

Electrical synapses enable circuits of coupled neurons to selectively respond to concurrent depolarizations of the membrane potential (Galarreta and Hestrin, 2001; Veruki and Hartveit, 2002; Curti et al., 2012; Alcami, 2018). Because a portion of the injected current flows into the electrically coupled neurons, electrical synapses act as current sinks, reducing the Rin of all neurons within the circuit. This reduction diminishes the efficacy of depolarizing inputs and their ability to elicit action potential firing (Getting and Willows, 1974; Bennett and Zukin, 2004; Alcami and Marty, 2013; Alcami, 2018; Davoine and Curti, 2019). Instead, synchronous inputs to all neurons in the circuit promote parallel variations in their membrane potentials, strongly reducing the voltage drop across gap junctions and thereby minimizing current leakage. In this way, synchronous synaptic inputs enhance changes in membrane potential, thereby facilitating neuronal activation. This property enables circuits of electrically coupled neurons to amplify the impact of coincident synaptic inputs while attenuating temporally dispersed ones, thereby promoting coincidence detection. This effect has been demonstrated both experimentally and theoretically (Smith and Vardi, 1995; Chillemi et al., 2007; Hjorth et al., 2009; Rabinowitch et al., 2013), highlighting the fact that coincidence detection emerges from the interplay between gap junction-mediated electrical coupling and intrinsic neuronal properties, with significant implications for the function of neural circuits (Rela and Szczupak, 2003; Curti et al., 2022).

### The IH current and its impact on electrical synaptic transmission

The role of various active membrane conductances in electrical synaptic transmission has been studied in the context of depolarizing changes in membrane potential, particularly concerning phenomena such as synchronization (Mann-Metzer and Yarom, 1999; Haas and Landisman, 2011), lateral excitation (Curti and Pereda, 2004; Trenholm et al., 2013), oscillations (Dugué et al., 2009; Trenholm et al., 2012) and coincidence detection (Davoine and Curti, 2019). In contrast, how transmission of hyperpolarizing signals is influenced by the intrinsic electrophysiological properties of electrically coupled neurons has received much less attention. In this regard, the IH current, a cationic current mediated by hyperpolarization-activated cyclic nucleotide-gated (HCN) channels, is widely expressed in neuronal populations, including those interconnected by electrical synapses. IH current is expressed in neurons of the inferior olive (Schweighofer et al., 1999; Devor and Yarom, 2002), the reticular nucleus of the thalamus (Landisman et al., 2002; Rateau and Ropert, 2006), pyramidal neurons of the hippocampus (Maccaferri et al., 1993; Mercer et al., 2006), cerebellar Golgi cells (Forti et al., 2006; Dugué et al., 2009), bipolar cells of the retina (Veruki and Hartveit, 2002; Müller et al., 2003), mitral cells of the olfactory bulb (Schoppa and Westbrook, 2002; Angelo and Margrie, 2011), neurons of the mesencephalic trigeminal nucleus (Khakh and Henderson, 1998; Curti et al., 2012) and molecular layer interneurons (MLI) of the cerebellar cortex (Mann-Metzer and Yarom, 1999; Southan et al., 2000), among others. The reversal potential of the IH current, located near the base of its activation curve, allows this conductance to act as a feedback mechanism that counterbalances changes in membrane potential. As a result, the IH current not only plays a role in establishing the resting membrane potential and Rin but it also modulates synaptic input integration, repetitive firing, and oscillatory activity (Pape, 1996; Biel et al., 2009; Benarroch, 2013). Furthermore, the widespread expression of the IH current in electrically coupled cells suggests that this conductance may critically influence the transmission of hyperpolarizing signals through electrical synapses.

In the retina, rod photoreceptors are interconnected through connexin36-containing gap junctions and express IH current (Sterling and Demb, 2004; Barrow and Wu, 2009). It has been shown that light-evoked currents induce long-lasting hyperpolarizations whose repolarizing phase is strongly shaped by the activation of HCN channels (Barrow and Wu, 2009). Remarkably, IH current activation imposes strong nonlinear characteristics to the spread of hyperpolarizing receptor potentials in the network of electrically coupled rods (Pang et al., 2024). In fact, the low-pass filter characteristics imposed by the passive properties of the cell membrane tend to introduce a time lag to the peak of voltage responses as they spread through a linear array of electrically coupled cells (Bennett and Zukin, 2004). This slowing effect on propagated hyperpolarizing signals makes them suitable to activate the slowly activating IH current in postsynaptic coupled rods. In turn, by counteracting membrane hyperpolarization, IH current activation results in a shortening of the time to peak responses in distant rods, thus compensating the time loss of signals traveling along the network of rods and facilitating synchronization of the output signals. As a result, the IH current might promote network synchrony of hyperpolarizing signals (Pang et al., 2024).

### Coupling strength and coincidence detection of hyperpolarizing inputs are strongly shaped by the IH current in MesV neurons

The specific voltage-dependent properties and activation kinetics of the IH current impart distinctive temporal dynamic characteristics to coupling strength. This phenomenon can be observed in pairs of electrically coupled mesencephalic trigeminal (MesV) neurons, which are coupled in small groups and exhibit a high density of IH current expression (Khakh and Henderson, 1998; Curti et al., 2012; Davoine and Curti, 2019). As shown in Figure 1Aa, injection of hyperpolarizing pulses of increasing intensity results in a progressively stronger activation of the IH current, as indicated by voltage membrane responses displaying increasing amplitude sags (Vm Cell 1). In turn, the postsynaptic coupled neuron shows near simultaneous voltage membrane deflections, although smaller in amplitude and of slower time course, resembling passive responses (Fig. 1Aa, Vm Cell 2). However, plots of the voltage response amplitude of the postsynaptic neuron as a function of that of the presynaptic neuron (whose slope corresponds to the coupling coefficient, CC), show that while this relationship is linear at the peak of hyperpolarizing responses (blue vertical dashed line in Fig. 1Aa and blue axis and symbols in Fig. 1Ab), it is sublinear at steady state (red vertical dashed line in Fig. 1Aa and red axis and symbols in Fig. 1Ab) for the larger-amplitude responses. In fact, when the pre- and postsynaptic responses are superimposed, after amplitude normalization, it can be readily appreciated that the coupling potential is more attenuated at steady state than at the transient hyperpolarizing response (peak) (Fig. 1Ba).

**Figure 1.**
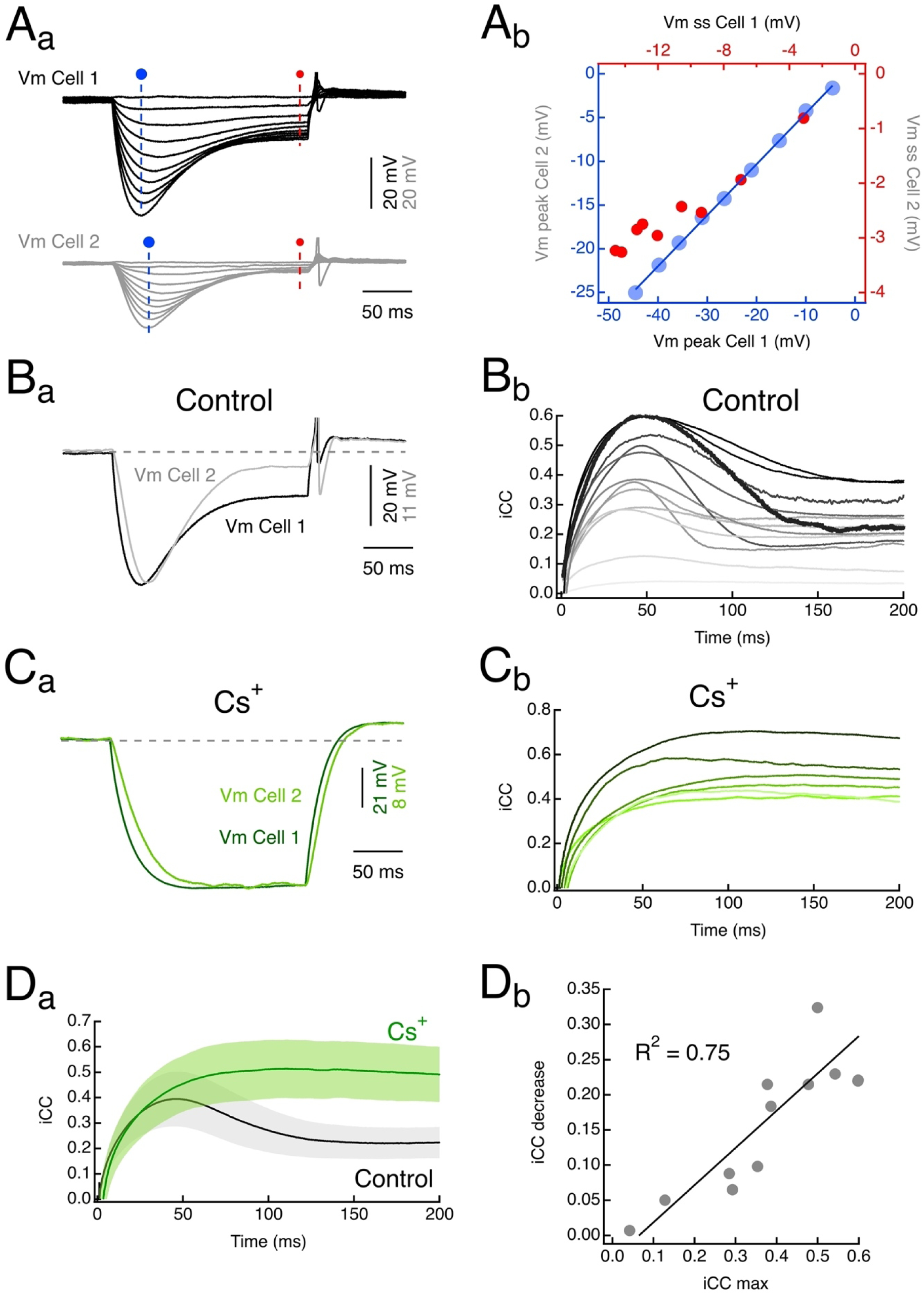
**(A_a_)** Paired recordings from electrically coupled MesV neurons. The injection of a series of hyperpolarizing current pulses (from -450 to 0 pA) of increasing intensity into one cell produces corresponding voltage responses in the same cell (Vm Cell 1) and in the coupled cell (Vm Cell 2). (**A_b_**) From the recordings in A_a_ the coupling strength was estimated by plotting the amplitude of membrane voltage changes in the postsynaptic cell (Vm Cell 2, ordinates) as a function of membrane voltage changes in the presynaptic cell (Vm Cell 1, abscissas), measured either at the peak of the hyperpolarizing responses (blue dashed line and circle in A_a_) or at steady state (red dashed line and circle in A_a_). While this relationship at the hyperpolarizing peak is linear in the whole range of tested membrane potentials, it becomes sublinear at steady state for the strongest hyperpolarizing pulses. (**B_a_**) Traces of the presynaptic (Vm Cell 1) and postsynaptic (Vm Cell 2) membrane voltage responses to a hyperpolarizing current pulse (-450 pA) in the presynaptic neuron, normalized to the peak of the responses. (**B_b_**) Plot of the instantaneous coupling coefficient (iCC) as a function of time during current pulse injection calculated from recordings like those depicted in B_a_ for 13 directions. Thicker trace represents example shown in B_a_. (**C_a_**) Superimposed traces of the presynaptic (Vm Cell 1) and postsynaptic (Vm Cell 2) membrane voltage responses to a hyperpolarizing current pulse (-400 pA) in the presence of Cs^+^ (5 mM) in the extracellular solution. (**C_b_**) Plot of the iCC as a function of time in the presence of Cs+ (2 – 5 mM) for 6 directions. (**D_a_**) Average iCC as a function of time calculated from plots in B_b_ and C_b_ in control conditions and in the presence of Cs^+^. Shaded areas represent 95% confidence intervals. (**D_a_**) Plot of the iCC reduction at steady state versus maximum value iCC for the directions depicted in B_b_ (circles). Data set was fitted with a straight-line function and the corresponding coefficient of determination is indicated. Data represent unpublished results.

Accordingly, the instantaneous coupling coefficient (iCC), calculated as the point-by-point ratio of the membrane voltage drop in the postsynaptic to the presynaptic cell, initially increases, reflecting the charging of the postsynaptic neuron’s membrane capacitance, and then undergoes a pronounced reduction from its peak value (Fig. 1Bb). In the pair shown in Figure 1Ba, the steady-state iCC declined to approximately half of its maximum value (black thick trace in Fig. 1Bb). Because the coupling strength is determined by both junctional resistance and the membrane resistance of the postsynaptic neuron (Curti and O’Brien, 2016), regulation of either parameter could contribute to the time-dependent reduction of the iCC. Interestingly, within this time window and range of transjunctional potentials, the resistance of these intercellular contacts does not show any sign of voltage dependence (Curti et al., 2012). This strongly suggests that the primary factor underlying the time-dependent decrease in iCC is the reduction in postsynaptic membrane resistance due to IH current activation, similarly to what has been determined in the context of the propagation of hyperpolarizing signals in the network of rod photoreceptors (see above) (Pang et al., 2024). Supporting this notion, blocking the IH current with extracellular Cs^+^ (2 - 5 mM) eliminates the IH current dependent sag (Fig. 1Ca) as well as the time-dependent decrease of the iCC in pairs of coupled MesV neurons (Fig. 1Cb and Da). The steady-state decrease of the iCC is significantly greater under control conditions compared to its reduction in the presence of Cs⁺. Specifically, iCC averaged 59.3 ± 14.8% (mean ± SD) under control conditions (n = 13 directions) and 94.7 ± 3.7% (mean ± SD) in the presence of Cs⁺ (n = 6 directions), relative to their respective maximum values. These values in control and in the presence of Cs^+^ are significantly different (P = 7.35 x 10^-3^, unpaired, two tailed Mann-Whitney Wilcoxon test). Notably, this time-dependent decrease in iCC is more pronounced in strongly coupled pairs than in weakly coupled ones, suggesting a dependence on the amplitude of the postsynaptic signals. Consistently, the magnitude of iCC reduction (defined as the difference between its maximum and steady-state values) exhibits a significant linear correlation with the maximum iCC (R^2^ = 0.75, linear correlation test p=0.00014, Fig. 1Db).

We hypothesize that this time-dependence of the iCC due to IH current activation may impact coincidence detection of hyperpolarizing inputs. To explore this possibility, negative current pulses were applied independently (Fig. 2A, left and middle) or simultaneously to pairs of electrically coupled MesV neurons (Fig. 2A, right). Hyperpolarizing peak responses during the simultaneous current injection are of bigger amplitude, as expected from the reduction of current leak through gap junctions in passive circuits. Interestingly, during the sag (corresponding to the activation of the IH current), simultaneous current injections evoke membrane voltage responses that become smaller in amplitude than those evoked by independent current injections (Fig. 2B and C). This suggests that by modulating coupling strength in a time-dependent manner, IH current activation can narrow the time window for coincidence detection of hyperpolarizing signals.

**Figure 2.**
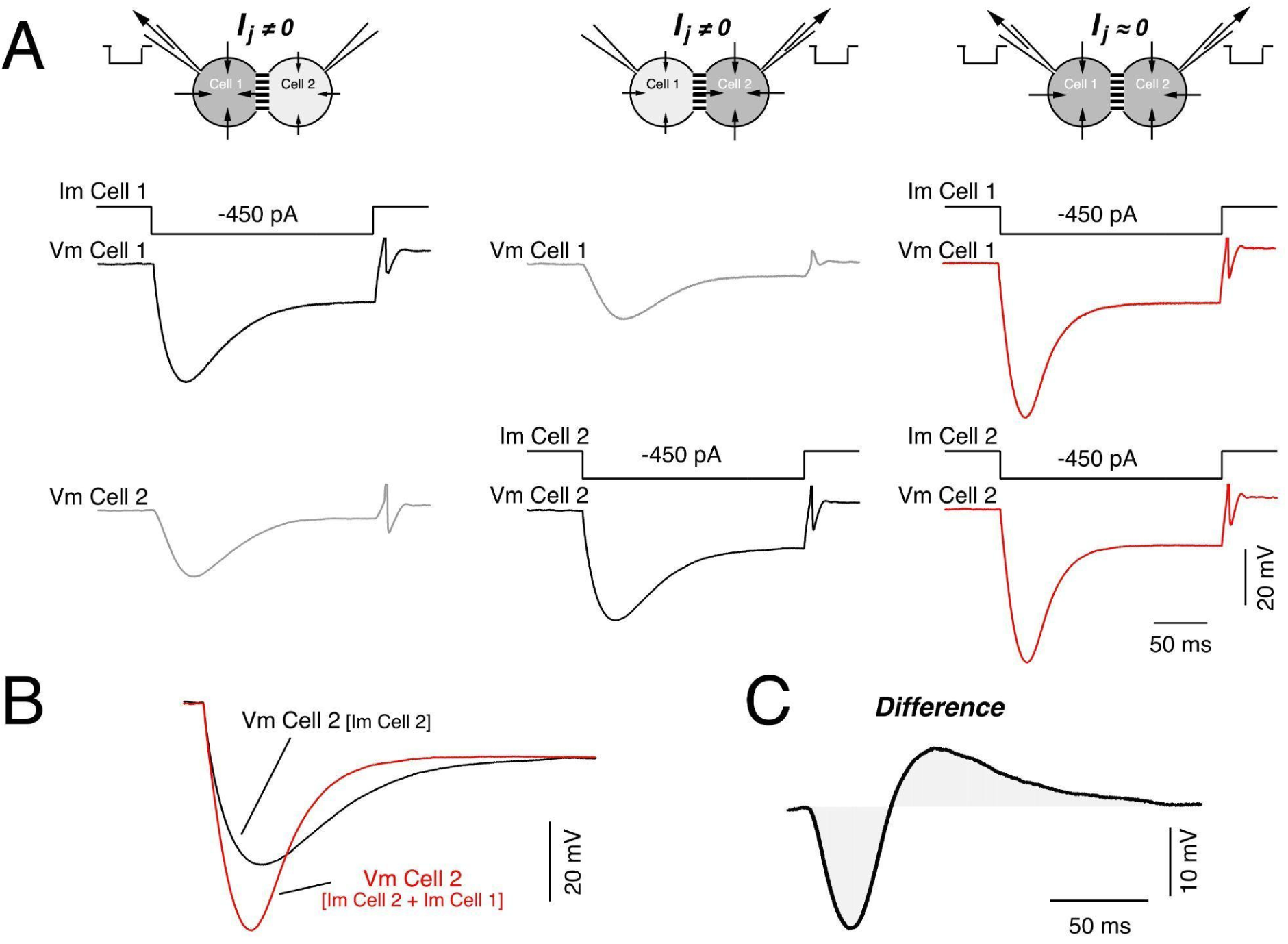
(**A**) Paired recordings from electrically coupled MesV neurons during the injection of a hyperpolarizing current pulse (Im) in cell 1 (left), in cell 2 (middle) or in both cells at the same time (right). Current pulses applied alternatively to each cell (left and middle traces) evoked the characteristic MesV neuron’s voltage membrane response along with their corresponding coupling potentials in the other cell due to the spread of part of the injected current through junctions (Ij ≠ 0). Current pulses of the same intensity applied simultaneously to both coupled neurons (left) induce larger responses due to the cancelation of the spread of current to the other cell (Ij ≈ 0). Schemes above each set of traces indicate the stimulation protocol and the corresponding current flow. (**B**) Recordings depicted in A of the membrane voltage responses of cell 2 when current pulses were injected only to this cell (Vm Cell 2 [Im Cell 2]) or simultaneously to both coupled cells (Vm Cell 2 [Im Cell 2 + Im Cell 1]) are illustrated superimposed to better appreciate the difference in amplitude and waveform. (**C**) Difference of the two recordings shown in B calculated as red minus black (the voltage response during simultaneous current injection minus the voltage response during current injection only to cell 2). Data represent unpublished results.

### Implications for the integration of chemical inhibitory synaptic inputs

Electrical transmission is usually evaluated in the hyperpolarized range at the steady state as a unique value of coupling strength. However, the present findings demonstrate that coupling is time-dependent and modulated by the IH current. Based on this we hypothesize that synchronous inhibitory inputs to groups of electrically coupled neurons might promote the activation of IH current by evoking hyperpolarizing responses which are larger in amplitude than voltage responses induced by temporally distributed inputs (coincidence detection). In turn, as IH current counteracts membrane hyperpolarization and dampens Rin, it dynamically sets the temporal window of the coincidence detection phenomenon.

Although MesV neurons provide a tractable model to examine the interaction of the IH current with hyperpolarizing inputs in a network of electrical coupled neurons, they lack inhibitory inputs, making it difficult to address its physiological significance. The contribution of IH current to electrical synapses-mediated coincidence detection of inhibitory inputs could nevertheless be further investigated in inhibitory interneuron networks. Indeed, interneurons typically express IH current and are interconnected by both electrical and chemical inhibitory synapses across the vertebrate brain. They include, among others, inhibitory interneurons in the hippocampus, neocortex, thalamus, and cerebellar cortex (Bartos et al., 2002; Hestrin and Galarreta, 2005; Aponte et al., 2006; Deleuze and Huguenard, 2006; Alcami and Marty, 2013; Rieubland et al., 2014a; Yang et al., 2018). Furthermore, electrical coupling has been described between principal cells as well, e.g. in the inferior olive, neocortex and hippocampus, which also express IH, suggesting that the coincidence detection of inhibitory inputs may also be modulated by the IH current in a number of principal cell networks (Llinás and Yarom, 1981; Draguhn et al., 1998; Wang et al., 2010).

We propose the circuits formed by cerebellar molecular layer interneurons (MLIs) as a model of choice for investigating the functional implications of the interaction between the IH current and coincidence detection of inhibitory inputs in electrically coupled networks. MLIs are interconnected through a combination of electrical and chemical inhibitory synapses (Llano and Gerschenfeld, 1993; Mann-Metzer and Yarom, 1999; Alcami and Marty, 2013; Kim et al., 2014; Rieubland et al., 2014b; Hoehne et al., 2020), and electrical coupling has been shown to mediate coincidence detection (Alcami, 2018). Additionally, MLIs express the IH current (Southan et al., 2000). Furthermore, MLI-mediated inhibition plays a crucial role in regulating the output of the cerebellum’s principal neuron, the Purkinje cell (Mittmann et al., 2005; Chu et al., 2012), likely influencing the temporal control of motor behavior and motor learning (Freeman, 2015).

### Molecular layer interneurons as a model for investigating the role of the interaction between electrical synapses and the IH current on inhibitory synaptic integration

MLIs are parvalbumin-positive, fast-spiking GABAergic interneurons that inhibit Purkinje cells (PCs) through both chemical synapses and ephaptic interactions, thereby regulating Purkinje cell firing frequency and rhythmic activity (Blot et al., 2016; Arlt and Häusser, 2020) and ultimately modulating cerebellar cortical output. They receive excitatory input from granule cells, the same source that excites PCs, forming a feedforward inhibitory circuit. This inhibitory control by the MLIs on the PCs and in particular on the firing frequency of the “simple” type of spikes is of utmost importance for fine motor coordination and the consolidation of the cerebellar-dependent motor learning^75^, such as the adaptation of the vestibulo-ocular reflex and classical conditioning like the eye blink reflex (Freeman, 2015)^76^.

As mentioned above, MLIs are interconnected via both electrical and chemical inhibitory synapses. They form strong GABAergic inhibitory connections onto each other, where a single presynaptic action potential can significantly delay the spontaneous firing of the postsynaptic MLIs (Lackey et al., 2024). In addition, the change in conductance associated with synaptic inhibition is capable of producing a marked drop in the Rin and the membrane time constant, modifying the cell’s intrinsic properties and therefore, neuronal integration (Häusser and Clark, 1997). Electrical coupling among MLIs is relatively strong (Mann-Metzer and Yarom, 1999; Alcami and Marty, 2013; Kim et al., 2014; Rieubland et al., 2014b; Hoehne et al., 2020) and has been reported to be more prevalent among specific subtypes of MLIs, which vary depending on the classification criterion, methods and preparation (Mann-Metzer and Yarom, 1999; Alcami and Marty, 2013; Kozareva et al., 2021).

Interestingly, recent transcriptomic analyses have identified two distinct subpopulations of MLIs (MLI1 and MLI2) based on the expression of GJD2, the gene encoding connexin36, the primary protein responsible for forming electrical synapses between mammalian neurons. Consistently, MLIs expressing the GJD2 gene (MLI1) are electrically coupled (Kozareva et al., 2021; Wang and Lefebvre, 2022; Lackey et al., 2024). Electrical synapses between MLIs promote their synchronization (Mann-Metzer and Yarom, 1999) and dynamically modify their firing probability and latency (Alcami, 2018). Indeed, coincidence detection mediated by electrical coupling enhances the firing of MLIs and reduces spike latency (Alcami, 2018), potentially contributing to the previously reported fast inhibition of PCs that outweighs excitation during sensory stimulation (Chu et al., 2012). Also, spikelets propagating through electrical synapses may induce net inhibition rather than excitation, narrowing the integration time window of EPSPs (Hoehne et al., 2020). Combined, the two modalities of synaptic transmission, electrical and chemical, play a central role in the output of cerebellar cortex circuits. While MLI1s primarily inhibit PCs, MLI2s are not electrically coupled and primarily inhibit MLI1s, suggesting that the effects of MLIs onto PCs excitability result from the contribution of two opposing mechanisms: on the one hand, direct inhibition from the electrically-coupled subtype (MLI1s), and on the other hand indirect disinhibition mediated by the inhibition from the other subtype (MLI2s) on MLI1s. Indeed, by inducing a firing pause of MLI1s, inhibitory inputs from MLI2s result in a desinhibition of PCs, and hence an increase of PC’s inhibitory influence in their postsynaptic target, the neurons in the deep cerebellar nuclei and various brainstem nuclei. These results raise the possibility that the efficacy of the inhibitory inputs onto the coupled MLI1 population may be critically determined by the degree of synchrony between these inputs and on the specific organization of projections from the MLI2s to MLI1s, particularly in terms of the degree of divergence.

Coincident inhibitory synaptic inputs onto electrically coupled MLIs are expected to produce a stronger inhibitory effect compared to independent ones. The resulting enhanced hyperpolarization from detection of coincident inputs may facilitate IH current activation, particularly during high-frequency repetitive synaptic activation, which promotes the summation of postsynaptic potentials. In turn, IH current activation may limit the duration of inhibition. MLIs express the IH current (Saitow and Konishi, 2000; Southan et al., 2000), supporting the notion that while simultaneous inhibitory inputs on electrically coupled MLIs are expected to induce a stronger control of their excitability, the time course of this effect is likely shaped by the dynamics of IH current activation. Thus, the integration of inhibitory inputs in the MLI network may be dynamically regulated by the activation of the IH current. Notably, MLIs express the HCN1 subunit (Luján et al., 2005), which has the fastest activation kinetics and the more positive voltage dependence of all the HCN subunits, rendering these channels particularly well-suited to interact with GABAergic synaptic inputs. In line with this, the time course for loading the membrane capacitance of coupled cells overlaps with the activation time scale of the IH current in MLIs (Alcami and Marty, 2013). While these aspects appear plausible, they remain to be experimentally investigated. Understanding the interplay between electrical and GABAergic synapses with the intrinsic properties, is crucial for elucidating how MLIs fine tune PCs activity. We propose that this neuronal circuit serves as an ideal model to test the functional implications of the interplay between electrical synapses-mediated coincidence detection and IH current on synaptic integration and network function.

### Concluding remarks and future directions

In this article, we have reviewed the role of electrical coupling and intrinsic properties, with a focus on the IH current, in regulating the temporal dynamics of hyperpolarizations, which we propose to be relevant for the integration of inhibitory synaptic potentials. Moreover, we characterized the time course of electrical synapse-mediated coincidence detection in MesV neurons and the contribution of the IH current in the hyperpolarizing range. Based on these results, we propose that cerebellar MLIs represent an optimal network for future studies on the impact of coincidence detection on synaptic input integration, a phenomenon likely to be broadly relevant in coupled networks receiving inhibitory synaptic inputs.

Temporal precision is an essential aspect in cerebellar physiology. In this regard, the interaction of these two modalities of synaptic transmission with the intrinsic neuronal properties, particularly the IH current, may influence the integration of inhibitory inputs by MLIs and hence the time window of their inhibitory effects on PCs. However, despite recent advances in the field, much remains to be investigated concerning the physiology of the MLI network. Another aspect that has remained largely unexplored is the effect of neuromodulatory systems on the physiology of MLIs. The IH current is under regulatory control by cyclic nucleotides levels (which in turn are under the regulatory control of several neuromodulatory substances) (Biel et al., 2009). By modulating the voltage-dependence of the IH current and the dynamics of its activation, it is expected that these modulatory systems exert a powerful control of MLI integration of synaptic inputs, and hence influence the timing of its inhibitory actions onto PCs. We hope that all of these questions will be answered in the years to come.

## Methods

Whole-cell patch-clamp recordings were performed on MesV neurons in current-clamp mode. Experimental procedures and solutions were identical to those described in our previous work (Curti et al., 2012). Briefly, transverse brainstem slices were obtained from Sprague-Dawley or Wistar rats (P7–P13) in accordance with the guidelines of the *Comisión Honoraria de Experimentación Animal* of the Universidad de la República (Uruguay), with measures taken to minimize the number of animals used. The coupling strength was assessed at room temperature in current-clamp by delivering hyperpolarizing current pulses with durations ranging from 200 to 400 ms into each cell alternatively. The resulting voltage deflections were recorded simultaneously in both the injected (presynaptic) and coupled (postsynaptic) cells. The instantaneous coupling coefficient (iCC) was determined as the point-by-point ratio of the membrane voltage change in the postsynaptic cell to that in the presynaptic cell, providing a zero-lag representation of coupling strength over time. In each pair of coupled neurons, the iCC values were estimated by this method in both directions and reported independently (two directions per coupled pair). During iCC calculation at the onset of current pulses, the slow change in membrane potential of the injected cell (presynaptic neuron) due to membrane capacitance charging led to near-zero denominators, resulting in artificially high iCC values. These values were removed. Data and statistical analyses were conducted using Igor Pro 7 (WaveMetrics).

## Acknowledgments

We thank Dr. Alain Marty for critical discussions and comments on an early version of this manuscript. This work was supported by a Comisión Sectorial de Investigación Científica (CSIC) of Universidad de la República, Uruguay, I+D grant to Sebastián Curti and Federico F. Trigo.

## Notes

### Competing Interest Statement

The authors have declared no competing interest.

